# Associations between childhood family emotional health, fronto-limbic grey matter volume, and saliva 5mC in young adulthood

**DOI:** 10.1101/2020.10.26.355347

**Authors:** JR Pfeiffer, Angela C. Bustamante, Grace S. Kim, Don Armstrong, Annchen R. Knodt, Karestan C. Koenen, Ahmad R. Hariri, Monica Uddin

## Abstract

**Background:** Poor family emotional health (FEH) during childhood is prevalent and impactful, and likely confers similar neurodevelopmental risks as other adverse social environments. Pointed FEH study efforts are underdeveloped, and the mechanisms by which poor FEH are biologically embedded are unclear. The current exploratory study examined whether variability in DNA methylation (DNAm) and fronto-limbic grey matter volume may represent pathways through which FEH may become biologically embedded.

**Results:** Self-reported childhood FEH was nominally associated with right hemisphere hippocampus (b=10.4, p=0.005), left hemisphere amygdala (b=5.3, p=0.009), and right hemisphere amygdala (b=5.8, p=0.016) volumes. Childhood FEH was also nominally associated with 49 DNAm MEs (p_range_=3×10^−6^ to 0.047). After limiting analyses to probes correlated between saliva and brain, saliva-derived DNAm MEs partially mediated the association between FEH and right hippocampal volume (Burlywood ME indirect effect b=-111, p=0.014), and fully mediated the FEH and right amygdala volume relationship (Pink4 ME indirect effect b=-48, p=0.026). Modules were enriched with probes falling in genes with immune, CNS, and metabolic functions.

**Conclusions:** Findings extend work highlighting neurodevelopmental variability associated with adverse social environment exposure during childhood by specifically implicating poor FEH, while informing a mechanism of biological embedding. FEH-associated epigenetic signatures could function as proxies of altered fronto-limbic grey matter volume associated with poor childhood FEH and inform further investigation into primarily affected tissues such as endocrine, immune, and CNS cell types.

## Background

Children can be exposed to an array of adverse social environments (ASEs) throughout their development. These include low socioeconomic status (SES), stressful life events (SLEs), trauma, and of particular interest, caregiver psychopathology. Caregiver psychopathology is prevalent in the United States; it is estimated that ~12.8 million parents suffer yearly from some form of mental illness (18.2%), and that ~2.7 million parents suffer yearly from a *serious* mental illness (3.8%)[1]. The psychological effects of living with mentally ill caregivers are notably deleterious. Children of caregivers with major depressive disorder (MDD), for example, experience more hostile, negative, and withdrawn parenting[2]. Estimates range from two to 13 times increased risk for children to develop either their caregiver’s mental illness or a mental illness different from their caregiver’s[3]. Children growing up in these conditions are also more likely to develop internalizing or externalizing behavioral problems, as well as social, cognitive, and academic difficulties[4, 5]. However, the mechanisms by which poor FEH are biologically embedded and produce these adverse outcomes are unclear.

The neuroimmune network hypothesis is one framework used to explain the physiological mechanisms via which ASEs and caregiver mental illness affect the mental health of offspring. The neuroimmune network hypothesis focuses on the integrated, bi-directional network of the central nervous system (CNS) and the immune system[6]. It posits that exposure to ASEs during childhood, an especially plastic window of development[7], impacts communication between peripheral inflammatory signals and brain regions responsible for threat, reward, executive control, memory, and adaptive behavioral/emotional responses (i.e. the fronto-limbic pathway), among others. Importantly, these functions are impaired in numerous mental illnesses, including but not limited to PTSD[8], MDD[9], anxiety disorders[10], bipolar disorder[11], and schizophrenia[11]. These inflammatory signals disrupt the inter-dependent functions of the front-limbic pathways, leading to altered behavioral states, and the pre-disposition to develop aberrant stress responses later in life[12]. These concepts are supported by a significant body of research that has shown immune system[13–15], hypothalamic-pituitary-adrenal (HPA)-axis[16, 17], and fronto-limbic pathway[17–21] associations with ASE exposure. More specifically, researchers have shown that childhood exposure to factors similar to poor FEH, such as maternal support and supportive/hostile parenting, are associated with lower hippocampus and amygdala grey matter volume later in life[22, 23]. Observed in association with ASE exposures, the signatures of morphometric variability within the fronto-limbic pathway are regarded as neural correlates of these exposures[17–23], and as neural endophenotypes of psychiatric illness[24–26].

The molecular mechanisms by which ASEs, including caregiver mental illness, become biologically embedded in the CNS are currently under investigation[27], and research has pointed to the importance of epigenetics, particularly 5’-methyl-cytosine (5mC) levels, in this process[28, 29]. 5mC serves as a mediator of gene by environment interaction[30–33], but it remains challenging to measure epigenetics in the living human brain -the primary etiologic tissue of interest in regards to mental health-related outcomes. This limitation has prompted investigation into epigenetic measures collected from peripheral tissue, such as saliva, which may serve as proxies for etiological tissue. Previous studies have provided a framework for the use of peripheral tissues in epigenomewide association studies (EWAS) and support the potential use of peripheral 5mC as a proxy for etiological tissue 5mC[34]. Further bolstering the notion that peripheral 5mC is an efficacious proxy for etiological tissue 5mC, is research showing that peripheral epigenetic measures can index changes in the HPA-axis[35, 36], immune system[37, 38], and the CNS[39–41]. However, these relationships do not directly indicate association between peripheral epigenetic measures and CNS-relevant endophenotypes of psychopathology. On this note, studies have used human structural and functional neuroimaging data in tandem with epigenetic measures but have primarily utilized candidate gene approaches. Measuring peripheral 5mC of the *SLC6A4*[42–44], *NR3C1[45,* 46], *FKBP5[47],* and *SKA2*[48, 49] genes, these studies have investigated associations between peripheral 5mC and variability in the structure and function of the frontal cortex, hippocampus, and amygdala. Findings suggest that locus-specific peripheral 5mC can index CNS structural alterations[42–49], and may statistically mediate ASE-induced CNS structural alterations[49].

Despite the evidence that peripheral 5mC can index CNS-related phenotypes, to date few studies, to our knowledge, have examined these relations in a hemisphere-specific manner within the brain. Importantly, numerous aspects of human behavior and biology are subject to hemisphere-specific brain lateralization[50–52]. This, coupled with evidence of hemisphere-specific fronto-limbic variability in association with ASEs in humans[17– 23], provide a solid framework to address the potential associations of poor FEH with *hemisphere-specific* volume measurements. Beyond the aforementioned reports, studies of poor FEH or caregiver mental illness on CNS structure are sparse and limited to biological offspring of parents with genetically heritable psychopathology, although they do investigate associations of exposure with outcome on a hemisphere-specific basis[53, 54]. These types of ASEs are also associated with changes in cell type-specific and tissue-specific 5mC[55]. However, to our knowledge, investigations into the role of poor FEH in association with neural endophenotypes of psychopathology development have yet to be reported, and therefore, the magnitude of risk associated with poor childhood FEH has not been elucidated. In addition, investigations into the potential epigenetic mechanisms explaining the biological embedding of poor FEH have yet to be carried out.

To address these gaps in the field, and to improve understanding of poor FEH exposure risk, the current exploratory study applied genome-scale approaches to assess whether saliva-derived DNA methylation (DNAm) measurements might index CNS endophenotypes of psychopathology in a sample of 98 young adult volunteers. We were specifically interested whether saliva-derived DNAm module eigengenes (MEs) might statistically mediate the relationship between poor FEH and hemisphere-specific fronto-limbic grey matter volume, while controlling for age, sex, cellular heterogeneity, genomic ancestry, past year SLEs, and total brain volume (TBV). Such a result may serve as a peripheral proxy of such CNS variability, while informing a *potential* biological mechanism of physiological embedding. Based on previous work, we hypothesized that identified 5mC modules would be enriched with CpG probes falling in genes with HPA-axis, immune system, and CNS-relevant gene ontology (GO) functions.

## Results

### Study participants

Descriptive statistics for demographic, psychosocial, and neuroimaging variables in study participants are shown in Table 1. FEH ranged from 34 to 70; the mean in the study sample was 60 (+/-8.5).

**Table 1.**
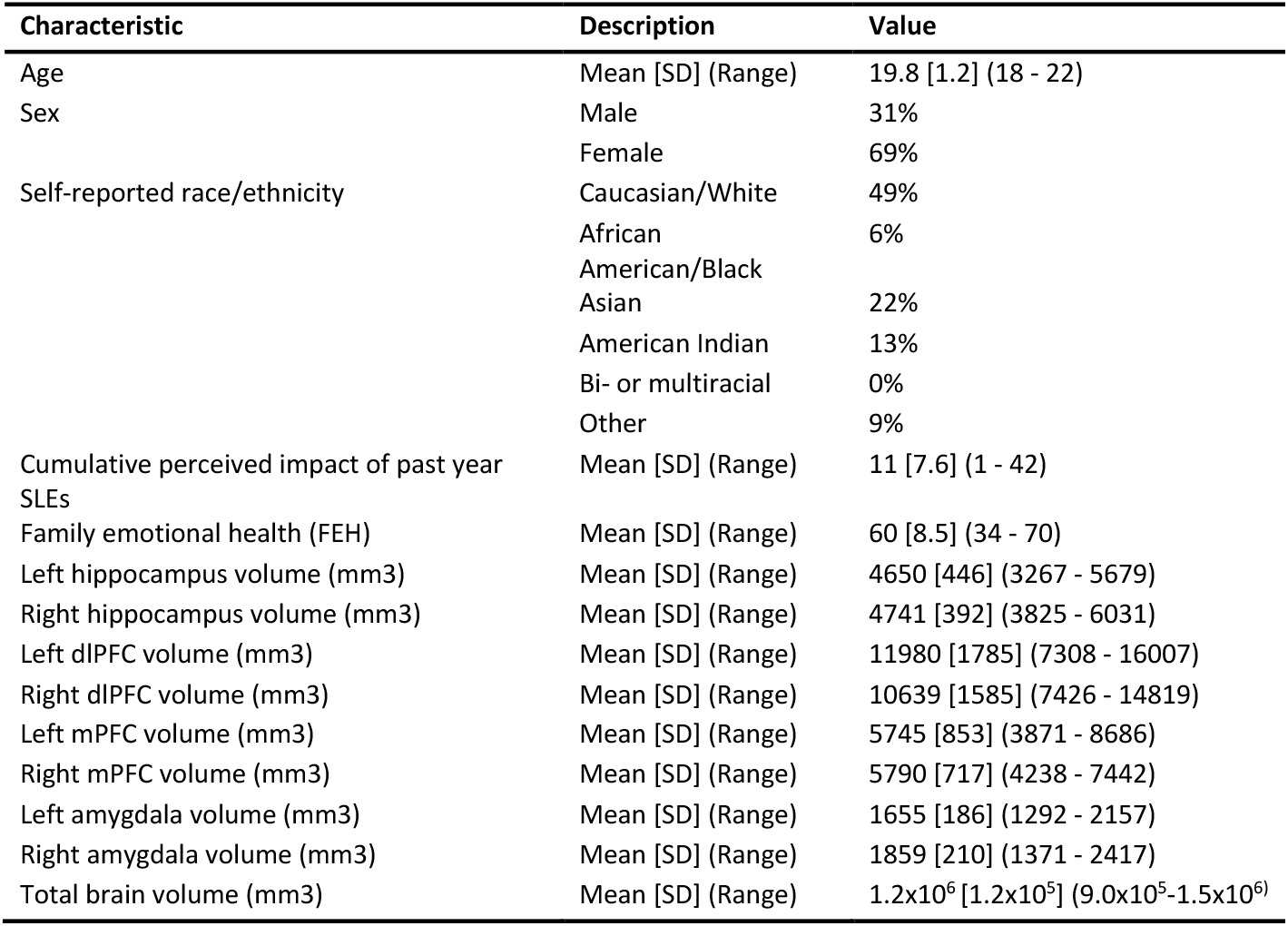
Demographic, psychosocial, and neuroimaging summary stats for the current sample (n = 98)

### Correlation analyses

Pearson correlations between variables used in the current study were mapped (Figure 2). Of note, a strong negative association was observed between FEH and past year SLEs (Pearson’s correlation: *r*=−0.44, p=7×10^−6^).

**Figure 1.**
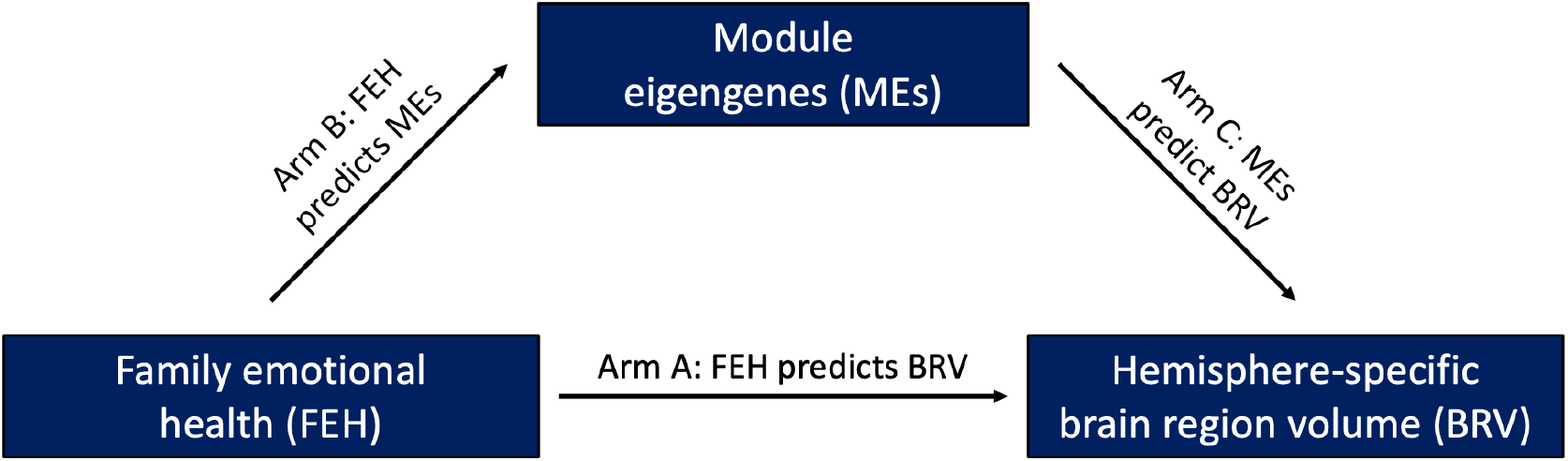
Conceptual model testing module eigengenes (MEs) as mediators of the hypothesized association between family emotional health (FEH) and variability in hemisphere-specific brain region volume (BRV). **Arm A.**FEH was used as a predictor of hemisphere-specific BRV, while including age, biological sex, four genomic ancestry MDS measures, past year SLEs, and total brain volume as covariates. **Arm B.** FEH was used as a predictor of ME value, while including past year SLEs as a covariate. The age, sex, and genomic ancestry effects on ME components were previously removed. **Arm C.** ME values were used as individual predictors of BRV, while including age, biological sex, four genomic ancestry MDS measures, past year SLEs, and total brain volume as covariates.

**Figure 2.**
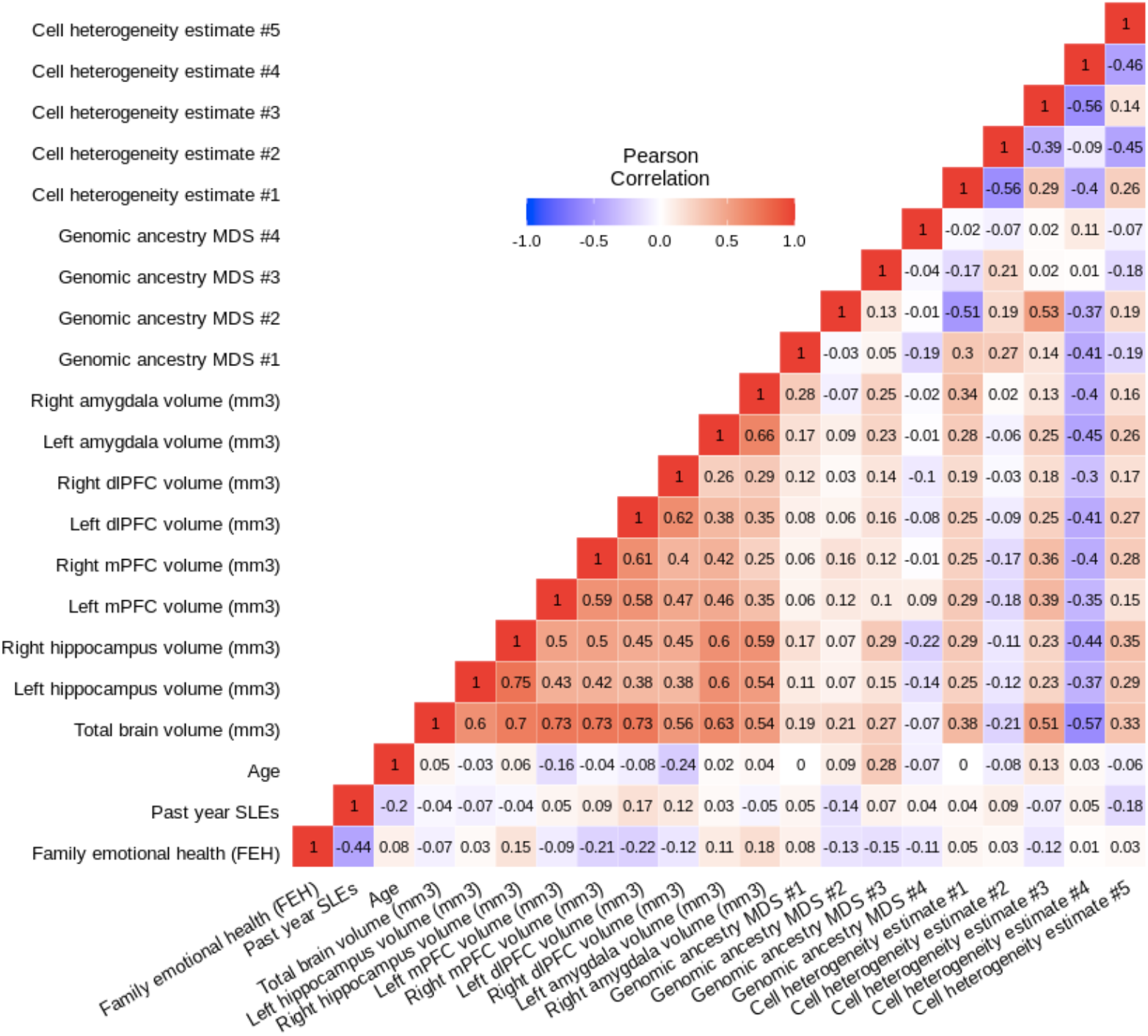
Pearson correlation heat map of variables used throughout the current analyses. **A.** A strong negative relationship was observed between FEH and past year SLEs (Pearson’s correlation: *r*=-0.44, p=7×10^−6^). Strong positive relationships are also observed between hemisphere-specific brain regions (Pearson’s correlation *r* range: 0.26 – 0.75).

### FEH predicts hemisphere-specific BRV

FEH was positively associated with right hippocampus (b=10.4, SE=3.6, t=2.9, p=0.005), left amygdala (b=5.3, SE=2.0, t=2.7, p=0.009), and right amygdala volumes (b=5.8, SE=2.3, t=2.4, p=0.016). These significant relationships were also observed in models without controlling for the covarying effect of TBV (right hippocampus p=0.015; left amygdala p=0.018; right amygdala p=0.023). FEH was not associated with left hippocampus (p=0.62), left dlPFC (p=0.10), right dlPFC (p=0.62), left mPFC (p=0.98), or right mPFC volume (p=0.09). In controlling for seventy-two tests at FDR=0.10, all three brain regions with nominal p<0.05 were BH-significant (Table 2). Regions associated with FEH were carried into following analyses.

**Table 2.** Family emotional health (FEH) predicts hemisphere-specific brain region volume (BRV)

|  |  |  |  |  |  |  |  |  |  |
| --- | --- | --- | --- | --- | --- | --- | --- | --- | --- |
| L. hipp vol. (mm3) | adj. R2 | RSE | b | SE | R. hipp vol. (mm3) | adj. R2 | RSE | b | SE |
| Model | 0.325*** | 369.2 |  |  | Model | 0.532*** | 255.9 |  |  |
| FEH |  |  | 2.6 | 5.2 | FEH |  |  | **10.4\*\*** | **3.6** |
| Sex (female) |  |  | -60.6 | 114.9 | Sex (female) |  |  | -88.5 | 79.7 |
| Age |  |  | -32.1 | 33.2 | Age |  |  | -8.4 | 23.0 |
| Cumulative impact of past year SLE |  |  | -2.5 | 5.9 | Cumulative impact of past year SLE |  |  | 4.1 | 4.1 |
| Total brain volume (mm3) |  |  | **0.002\*\*\*** | **0.0005** | Total brain volume (mm3) |  |  | **0.002\*\*\*** | **0.0003** |
| L. dlPFC vol. (mm3) | adj. R2 | RSE | b | SE | R. dlPFC vol. (mm3) | adj. R2 | RSE | b | SE |
| Model | 0.564*** | 1182.0 |  |  | Model | 0.344** | 1254.0 |  |  |
| FEH |  |  | -27.9 | 16.7 | FEH |  |  | -8.8 | 17.7 |
| Sex (female) |  |  | 366.8 | 367.9 | Sex (female) |  |  | 219.2 | 390.3 |
| Age |  |  | -82.5 | 106.3 | Age |  |  | **-323.2\*\*** | **112.8** |
| Cumulative impact of past year SLE |  |  | 28.3 | 18.9 | Cumulative impact of past year SLE |  |  | 10.9 | 20.0 |
| Total brain volume (mm3) |  |  | **0.013\*\*\*** | **0.0015** | Total brain volume (mm3) |  |  | **0.008\*\*\*** | **0.0016** |
| L. mPFC vol. (mm3) | adj. R2 | RSE | b | SE | R. mPFC vol. (mm3) | adj. R2 | RSE | b | SE |
| Model | 0.559*** | 557.6 |  |  | Model | 0.539*** | 479.1 |  |  |
| FEH |  |  | 0.2 | 7.9 | FEH |  |  | -11.6 | 6.8 |
| Sex (female) |  |  | 159.9 | 173.6 | Sex (female) |  |  | 53.1 | 149.1 |
| Age |  |  | -102.9* | 50.2 | Age |  |  | -11.6 | 43.1 |
| Cumulative impact of past year SLE |  |  | 4.8 | 8.9 | Cumulative impact of past year SLE |  |  | 5.7 | 7.7 |
| Total brain volume (mm3) |  |  | **0.006\*\*\*** | **0.0007** | Total brain volume (mm3) |  |  | **0.005\*\*\*** | **0.0006** |
| L. amygdala vol. (mm3) | adj. R2 | RSE | b | SE | R. amygdala vol. (mm3) | adj. R2 | RSE | b | SE |
| Model | 0.404*** | 139.2 |  |  | Model | 0.356*** | 166.2 |  |  |
| FEH |  |  | **5.2\*\*** | **2.0** | FEH |  |  | **5.8\*** | **2.3** |
| Sex (female) |  |  | -34.5 | 43.3 | Sex (female) |  |  | -34.3 | 51.7 |
| Age |  |  | -3.6 | 12.5 | Age |  |  | -5.1 | 15.0 |
| Cumulative impact of past year SLE |  |  | 3.5 | 2.2 | Cumulative impact of past year SLE |  |  | 0.7 | 2.7 |
| Total brain volume (mm3) |  |  | **0.001\*\*\*** | **0.0002** | Total brain volume (mm3) |  |  | **0.001\*\*\*** | **0.0002** |
| *** P < 0.001, ** P < 0.01, * P < 0.05, P: BH-significant (bolded) |  |  |  |  |  |  |  |  |  |
| Four genomic ancestry multi-dimensional scaling measures were included in each model, but are not shown here. |  |  |  |  |  |  |  |  |  |
| RSE: relative standard error, adj: adjusted, b: estimate, SE: standard error, L: left, R: right, hipp: hippocampus, vol: volume |  |  |  |  |  |  |  |  |  |

### FEHpredicts ME values

FEH was associated with 49 MEs (b_min_=−0.006, b_max_=0.006, p_min_=3×10^−6^, p_max_=0.047). Twenty-nine out of 49 MEs achieved BH-significance, including the Burlywood and Pink4 MEs, taking 194 tests into account at FDR=0.10 (Supplementary Table 2).

### ME values predict hemisphere-specific BRV

Forty-nine MEs nominally associated with FEH were tested for association with right hippocampus, left amygdala, and right amygdala volumes (Supplementary Table 3). Seven MEs were nominally associated with right hippocampus volume, four of which were BH-significant: Burlywood (b=874.2, SE=252.1, t=3.5, p=8×10^−4^) (Figure 3a), Darkolivegreen1 (b=770.0, SE=258.6, t=3.0, p=0.004), Thistle2 (b=728.1, SE=261.8, t=2.8, p=0.007), and Chocolate2 (b=-713.3, SE=259.3, t=-2.8, p=0.007). The Darkgray ME (b=-374.2, SE=140.1, t=-2.7, p=0.009) (Figure 3b) was negatively associated with left amygdala volume, in addition to the Darkolivegreen ME (b=-300.6, SE=142.2, t=-2.1, p=0.037). The Lavenderblush2 ME was positively associated with left amygdala volume (b=295.1, SE=144.5, t=2.0, p=0.044). Finally, the Pink4 ME (b=467.5, SE=165.8, t=2.8, p=0.006) (Figure 3c) was positively associated with right amygdala volume. In controlling for 49 tests within each of the three BRVs at FDR=0.10, only the aforementioned MEs associated with right hippocampus volume were BH-significant.

**Figure 3.**
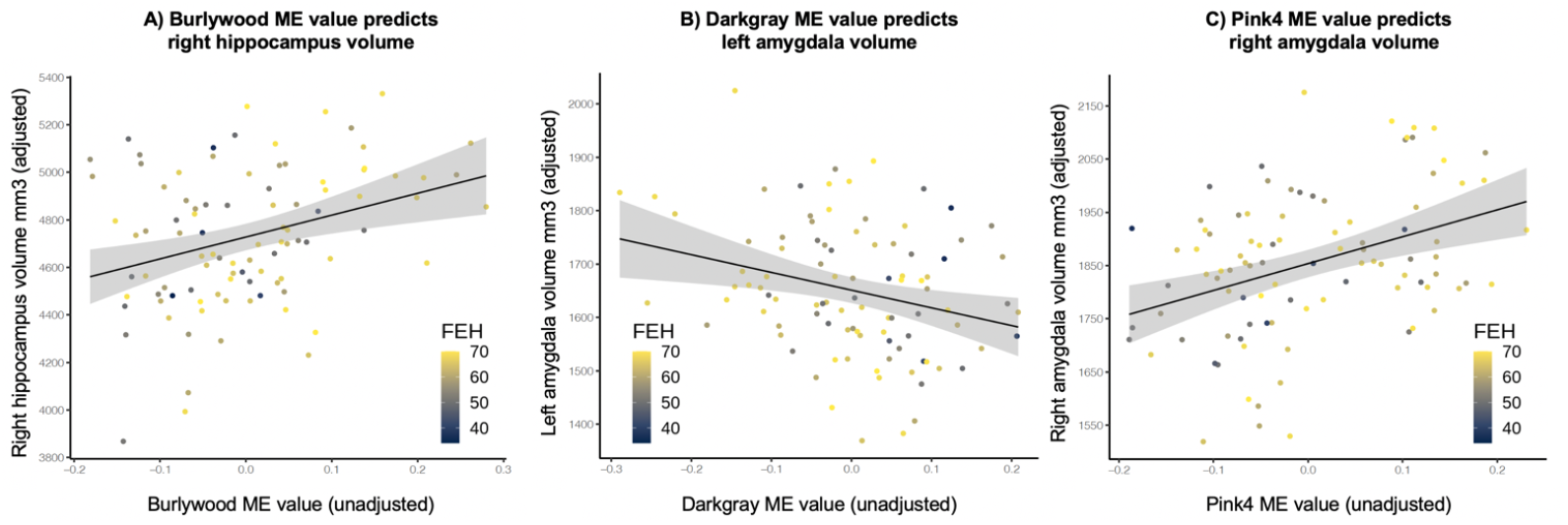
ME values are associated with high right hippocampal, low left amygdala, and high right amygdala volume. BRVs values shown are adjusted by covariates. Covariates across all models: age, sex, four genomic ancestry measures, past year SLEs, and total brain volume. The line of best fit (via least squares) is shown with a grey 95% SE confidence range. **A.** High Burlywood ME value is associated with high right hippocampal volume (b=874.2, SE=252.1, t=3.5, p=0.0008). **B.** High Darkgray ME value is associated with low left amygdala volume (b=- 374.2, SE=140.1, t= −2.7, p=0.009). **C.** High Pink4 ME value is associated with high right amygdala volume (b=467.5, SE=165.8, t=2.8, p=0.006).

### ME mediation

Eleven MEs were tested for mediation between FEH and BRVs. The Burlywood ME was a partial *statistical* mediator between FEH and right hippocampus volume (b_TE_=−366, p=8×10^−4^; b_IDE_=−111, p=0.014; b_DE_=−254, p=0.037). The TE indicated that right hippocampal volume was 366 mm^3^ less under poor FEH conditions compared to high FEH conditions, while the IDE of the Burlywood ME was accountable for 111 mm^3^ (30%) of that effect. Without controlling for the covarying effect of TBV, the Burlywood ME was a full mediator (b_TE_=−376, p=0.006; b_IDE_=−114, p=0.031; b_DE_=−261, p=0.071). The Darkolivegreen1 (b_TE_=−369, p=0.001; b_IDE_=−66, p=0.042; b_DE_=−303, p=0.008) and Thistle2 (b_TE_=−373, p=0.002; b_IDE_=−64, p=0.025; b_DE_=−309, p=0.010) MEs were also partial statistical mediators of the FEH and right hippocampus volume relationship. The Thistle ME was also a partial mediator in analyses without controlling for TBV (b_TE_=−382, p=0.007; b_IDE_=−85, p=0.017; b_DE_=−297, p=0.042). On the other hand, the Chocolate2, Cornflowerblue, Aliceblue, and Yellow MEs were neither partial nor full mediators of the relationship (p_TE_<0.05; p_IDE_>0.05; p_DE_<0.05).

None of the Darkgray (b_TE_=-183, p=0.014; b_IDE_=−47, p=0.095; b_DE_=−135, p=0.094), Darkolivegreen (b_TE_=−185, p=0.013; b_IDE_=−32, p=0.205; b_DE_=−153, p=0.057), or Lavenderblush2 (b_TE_=−187, p=0.011; b_IDE_=−30, p=0.181; b_DE_=−156, p=0.044) MEs were mediators of the relationship between FEH and left amygdala volume. However, the significant TE values indicated ~185 mm^3^ lower left amygdala volume in poor FEH conditions. Regarding FEH and right amygdala volume, Pink4 ME value was a full statistical mediator of the relationship (b_TE_=−204, p=0.017; b_IDE_=−48, p=0.026; b_DE_=−156, p=0.069), indicating that right amygdala volume was 204 mm^3^ less in poor FEH conditions than in high FEH conditions. Results additionally indicate that Pink4 ME value accounted for 48 mm^3^ (24%) of the aforementioned effect. Without controlling for the statistical effect of TBV, the Pink4 ME was again a full mediator of the FEH and right amygdala volume relationship (b_TE_=−208, p=0.017; b_IDE_=−52, p=0.025; b_DE_=−157, p=0.087). In controlling for 33 tests at FDR=0.10, all nominally significant ME IDE’s, DE’s, and TE’s were BH-significant (Table 3). Mediation analyses were then performed on individual probes from the Pink4 module in order to assess locus-specific effects.

**Table 3.** Module eigengenes (MEs) mediating observed family emotional health (FEH) and brain region volume (BRV) relationships

|  |  |  |  |
| --- | --- | --- | --- |
| <u>Burlywood: Right hippocampus</u> | <u>b</u> | <u>95% CI Lower</u> | <u>95% CI Upper</u> |
| Average indirect effect (IDE) | **-111.2\*** | -255.5 | -17.9 |
| Average direct effect (DE) | **-254.4\*** | -528.2 | -15.8 |
| Average total effect (TE) | **-365.7\*\*\*** | -641.0 | -151.0 |
| <u>Darkolivegreen1: Right hippocampus</u> | <u>b</u> | <u>95% CI Lower</u> | <u>95% CI Upper</u> |
| Average indirect effect (IDE) | **-65.6\*** | -161.3 | -1.6 |
| Average direct effect (DE) | **-302.9\*\*** | -553.6 | -84.3 |
| Average total effect (TE) | **-368.5\*\*** | -623.6 | -152.7 |
| <u>Thistle2: Right hippocampus</u> | <u>b</u> | <u>95% CI Lower</u> | <u>95% CI Upper</u> |
| Average indirect effect (IDE) | **-64.4\*** | -156.6 | -5.6 |
| Average direct effect (DE) | **-308.5\*** | -567.7 | -79.2 |
| Average total effect (TE) | **-372.9\*\*** | -640.1 | -150.5 |
| <u>Chocolate2: Right hippocampus</u> | <u>b</u> | <u>95% CI Lower</u> | <u>95% CI Upper</u> |
| Average indirect effect (IDE) | -82.8 | -204.7 | 4.9 |
| Average direct effect (DE) | **-281.5\*** | -585.5 | -24.2 |
| Average total effect (TE) | **-364.3\*\*** | -635.0 | -144.5 |
| <u>Cornflowerblue: Right hippocampus</u> | <u>b</u> | <u>95% CI Lower</u> | <u>95% CI Upper</u> |
| Average indirect effect (IDE) | -48.3 | -132.5 | 14.3 |
| Average direct effect (DE) | **-321.5\*\*** | -606.7 | -96.5 |
| Average total effect (TE) | **-369.8\*\*** | -641.6 | -151.8 |
| <u>Aliceblue: Right hippocampus</u> | <u>b</u> | <u>95% CI Lower</u> | <u>95% CI Upper</u> |
| Average indirect effect (IDE) | -53.4 | -146.6 | 6.6 |
| Average direct effect (DE) | **-317.6\*\*** | -591.1 | -98.8 |
| Average total effect (TE) | **-371.1\*\*\*** | -642.8 | -162.5 |
| <u>Yellow: Right hippocampus</u> | <u>b</u> | <u>95% CI Lower</u> | <u>95% CI Upper</u> |
| Average indirect effect (IDE) | -39.7 | -115.2 | 14.7 |
| Average direct effect (DE) | **-332.2\*\*** | -610.1 | -113.6 |
| Average total effect (TE) | **-371.9\*\*** | -642.0 | -158.7 |
| <u>Darkgray: Left amygdala</u> | <u>b</u> | <u>95% CI Lower</u> | <u>95% CI Upper</u> |
| Average indirect effect (IDE) | -47.4 | -115.8 | 8.9 |
| Average direct effect (DE) | -135.2 | -293.1 | 22.0 |
| Average total effect (TE) | **-182.5\*** | -324.2 | -38.6 |
| <u>Darkolivegreen: Left amygdala</u> | <u>b</u> | <u>95% CI Lower</u> | <u>95% CI Upper</u> |
| Average indirect effect (IDE) | -32.1 | -89.7 | 21.8 |
| Average direct effect (DE) | -152.5 | -310.8 | 4.6 |
| Average total effect (TE) | **-184.6\*** | -323.9 | -39.9 |
| <u>Lavenderblush2: Left amygdala</u> | <u>b</u> | <u>95% CI Lower</u> | <u>95% CI Upper</u> |
| Average indirect effect (IDE) | -30.4 | -93.4 | 13.7 |
| Average direct effect (DE) | **-156.4\*** | -301.0 | -3.0 |
| Average total effect (TE) | **-186.9\*** | -324.4 | -44.0 |
| <u>Pink4: Right amygdala</u> | <u>b</u> | <u>95% CI Lower</u> | <u>95% CI Upper</u> |
| Average indirect effect (IDE) | **-47.9\*** | -117.4 | -3.7 |
| Average direct effect (DE) | -156.1 | -338.9 | 12.9 |
| Average total effect (TE) | **-204.1\*** | -380.1 | -44.2 |
| *** P < 0.001, ** P < 0.01, * P < 0.05, P: BH-significant (bolded) |  |  |  |
| ME: module eigenvalue, b: estimate, CI: confidence interval |  |  |  |

### Probe-wise mediation

Three out of 21 probes from the Pink4 module were full mediators between FEH and right amygdala volume: cg22325292 (b_TE_=−204, p=0.013; b_IDE_=−53, p=0.018; b_DE_=−151, p=0.087), cg02398342 (b_TE_=− 204, p=0.014; b_IDE_=−44, p=0.038; b_DE_=−161, p=0.060), and cg00809820 (b_TE_=−205, p=0.013; b_IDE_=−48, p=0.049; b_DE_=−157, p=0.064). These three probes also had extremely high Pearson correlation values with the Pink4 ME (*r*>0.93, p<2×10^−44^), indicating that they are strong representatives of the Pink4 ME. In controlling for 63 tests at FDR=0.10, all nominally significant probe IDE’s, DE’s, and TE’s were BH-significant (Supplementary Table 4).

### Gene set enrichment analysis

We performed GSEA using probe M-values as predictors of FEH and used resultant p-values to facilitate the testing of 3,186 GO-terms. After redundancy reduction, 45 BH-significant GO-terms remained for interpretation. CNS-related GO-terms included: beta-amyloid clearance (GO:0097242, p=8×10^−11^, rank=2), filopodia assembly (GO:0046847, p=2×10^−8^, rank=5), catecholamine metabolic process (GO:0006584, p=4×10^−5^, rank=11), and positive regulation of neuron apoptotic process (GO:0043525, p=0.013, rank=25) among others. Although immune-related terms were limited, one was present in the top three: cytokine receptor activity (GO:0004896, p=8×10^−11^, rank=3). Numerous metabolic functions were identified: negative regulation of stress-activated MAPK cascade (GO:0032873, p=4×10^−12^, rank=1), NAD metabolic process (GO:0019674, p=4×10^−9^, rank=4), and TOR signaling (GO:0031929, p=3×10^−4^, rank=12) among others. A complete list of BH-significant GO-terms can be found in Supplementary Table 5.

## Discussion

The current exploratory study examined whether variability in DNAm and fronto-limbic grey matter volume represent pathways through which FEH becomes biologically embedded. Based on previous work, we hypothesized that 5mC modules would be enriched for immune system[13–15], HPA-axis[16, 17], and CNS-relevant[17–21] GO-terms. Our study findings indicated that exposure to poor FEH during childhood was associated with CNS endophenotypes of psychiatric illness, and that a subset of saliva-derived 5mC measurements *statistically* mediated this relationship. Additionally, we found the mediating 5mC modules were enriched with probes in genes with CNS-relevant and immune system GO-terms. Finally, we found that the underlying FEH-associated methylomic network was enriched with CNS-related, immune system, and metabolic gene sets. Overall, we posit that the FEH-associated epigenetic signatures could function as proxies of altered fronto-limbic grey matter volume associated with poor childhood FEH; peripheral epigenetic signatures indexing our relationships of interest may be explained by peripheral inflammation related to development of stress-related psychopathology, thereby supporting the neuroimmune network hypothesis[6].

The relationships observed between poor childhood FEH and left/right amygdala volume in the current study mirrored relationships observed throughout the literature regarding direction of effect and magnitude, but not hemisphere-specificity[17, 18]. Studies show hemisphere-specific effects of ASEs on amygdala volume, with stressors exerting notable statistical effects on left but not right amygdala volume. In one such prospective longitudinal study, SLEs negatively predicted left, but not right, amygdala volume in children with low to average polygenic risk scores. They showed that children exposed to the highest level of SLEs had ~9% less left amygdala volume than those exposed to the lowest levels of SLEs[17]. A more recent study showed lower left amygdala volumes in children who had experienced early neglect, low SES, or physical abuse compared to non-exposed controls[18]. Although we observed bilateral amygdala grey matter volume associations with poor childhood FEH exposure, our study did show a similar magnitude of effect; poor childhood FEH exposure was explanatory (DE) of - 8.9% difference in left and −8.4% difference in right amygdala volume. The peripheral 5mC signature (Pink4 ME) mediating right amygdala volume and FEH accounted for −2.5% of additional volumetric difference (IDE).

Similar to our amygdala-related findings, the reported relationship between poor childhood FEH and low hippocampus volume supports previous findings from the field regarding direction and estimated magnitude of effect, but not hemisphere-specificity[22, 23]. In a prospective longitudinal study, researchers focused on childhood “maternal support” as their exposure of interest, finding that maternal support of children, three to five years old, was associated with increased hippocampal volume in both hemispheres later in childhood (seven to thirteen years old). Specifically, they found that children exposed to low maternal support during that time span had a difference in hippocampal volume of −7.1%[22]. This magnitude closely mirrors the findings of the current study, which show poor childhood FEH has a DE that explains −6.1% difference in right hippocampal volume, and peripheral 5mC signatures have an IDE responsible for an additional −1.7% of difference. A more recent study from the same group found that the positive association between SES and hippocampal volume was mediated by “supportive/hostile parenting” in both hemispheres, but only by SLEs in left hippocampus[23]. These studies identified significant associations of maternal support and supportive/hostile parenting in *both* hippocampal hemispheres, whereas the current study identified a significant association only in right hippocampus.

No salient effects of FEH were observed in dlPFC or mPFC, in either hemisphere. This finding does not support research showing deleterious effects of ASEs on frontal cortex morphometry[56–58]. Our findings across fronto-limbic brain regions imply that poor childhood FEH has specific morphometric associations with subcortical structures responsible for memory, avoidance, fear, stress, and negative valence, but not cortical structures managing those functions.

Beyond the observed associations between poor childhood FEH and fronto-limbic brain morphometry, we were interested in the peripheral epigenetic signatures that index the relationships, and that provide a potential mechanism of biological embedding of ASEs. The Pink4 module, which fully mediated the relationship between poor childhood FEH and right amygdala volume in both TBV-controlled and non TBV-controlled models, is composed of 21 probes mostly mapped to known genes *(SNORD123, TBCD, FN3K, NRXN3, GLB1L2, SBF2, PSMB1, SYT1, BEST2, TBATA,* and *GNA12).* GO-terms associated with mapped genes include GO:0048487 beta-tubulin binding, GO:0038023 signaling receptor activity, GO:0019905 syntaxin binding, and GO:0031683 G-protein beta/gamma-subunit complex binding. Three out of 21 probes from the Pink4 module were full mediators of FEH and right amygdala volume: cg22325292, cg02398342, and cg00809820. Probes cg22325292 and cg02398342 exist in the sixth of six exons of the *FN3K* gene and fall in a putative CpG island and DNaseI hypersensitive region ~1,000 base pairs upstream of the *TBCD* transcription start site (TSS)[59, 60]. The main *TBCD* protein isomer plays a major role in the assembly of microtubules[61], the cell-cycle progression to mitosis[62], and neuronal morphogenesis[63]. Hypermethylation of the *TBCD* gene in CD4+ T-cells is also associated with rheumatoid arthritis[64], an autoimmune disorder associated with stress exposure[65]. Additionally, cg02398342 falls in the transcription factor binding site of the *EGR1* protein, which has integral, dynamic interactions with genes responsible for vesicular release and endocytosis, neurotransmitter metabolism and receptors, and actin cytoskeleton organization[66]. These interactions facilitate *EGR1’s* significant impact on synaptic and neuronal activation. Results suggest that the association of poor childhood FEH with right amygdala volume is indexed and *statistically* mediated by peripheral epigenetic signatures relevant to synapse development and cytoskeleton organization.

Three modules were partial mediators of the relationship between right hippocampus volume and poor childhood FEH: Burlywood, Darkolivegreen1, and Thistle2. The Burlywood module was a full mediator in the non TBV-controlled mediation model, implying that this peripheral epigenetic signature exerts more statistical effect on absolute right hippocampus volume, agnostic of TBV, through poor childhood FEH. Six of the 11 Burlywood probes are mapped to known genes *(MSH2, ATXN7L1, ODF2, SLC22A6, TGFB3,* and *DYX1C1)* with GO-terms including GO:0005245 voltage-gated calcium channel activity, GO:0002700 regulation of production of molecular mediator of immune response, and GO:0043524 negative regulation of neuron apoptotic process. GO-terms associated with probes from the Darkolivegreen1 and Thistle2 modules include GO:0001829 trophectodermal cell differentiation, GO:0045087 innate immune response, GO:0042552 myelination, and GO:0010506 regulation of autophagy, among others[59, 60]. Results imply that the associations of poor childhood FEH with right hippocampal volume are indexed by peripheral epigenetics signatures related to immune response and CNS cell development/lifecycle.

The top three GO-terms from our methylome network analysis were: 1. negative regulation of stress-activated MAPK cascade 2. beta-amyloid clearance 3. cytokine receptor activity. The MAPK cascade has long been established as a key driver of eukaryotic signal transduction, but more recently as an integral contributor to cell proliferation, differentiation, and inflammatory processes[67]. There is also a building body of evidence suggesting a significant role of the MAPK cascade in mental health outcomes. In a mouse model, modulation of the MAPK cascade in the forebrain is associated with both anxiety-like and depressive-like behaviors[68]. When p38 MAPK protein is selectively knocked out (KO) of the dorsal raphe nucleus, rodents subjected to social defeat stress show significantly reduced social avoidance compared to wild-type animals[69]. Additionally, pro-inflammatory cytokine administration induces a state of increased serotonergic CNS activity (canonically thought to be depleted in MDD), and induction towards that state is blocked with p38 MAPK inhibition[70]. In humans, MDD is a common comorbidity of rheumatoid arthritis (RA)[71]; peripheral inflammation is a hallmark of RA and is also observed in MDD patients[72]. Therefore, it is hypothesized that within the context of psychopathology development, environmental stressors induce peripheral cytokine signaling that communicates with fronto-limbic brain regions including the amygdala, hippocampus, and frontal cortex through mechanisms including the MAPK cascade[73]. To this end, numerous RA and anti-depressant drugs are observed to reduce canonical disease symptoms, while also reducing clinical inflammation markers and MAPK signaling[73].

It appears, then, that variability in peripheral DNAm and fronto-limbic grey matter volume represent pathways through which FEH becomes biologically embedded, with DNAm signatures that mediate the relation between FEH and grey matter volume being especially enriched with GO-terms related to the peripheral inflammatory sequela of stress-related psychopathology development. To our knowledge, the degree to which peripheral 5mC serves as a statistical mediator between poor childhood FEH (or ASEs in general) and variable fronto-limbic brain morphometry had not been previously elucidated. In addition, the observed GO-terms support potential mechanisms of biological embedding that are actively being considered in the field[68–70, 73, 74].

Dimension reduction techniques used throughout our research represent the foremost strengths of this study. These methods focus the analysis onto loci with greater prospect for proxy or surrogate status with etiologically relevant CNS tissue, and reduce the burden of multiple hypothesis testing. Clustering similarly methylated probes creates a relatively small number of modules which potentially contain probes from functionally related genes. On the other hand, limitations of the current study include relatively small sample size, lack of replication in an independent cohort, the balance of biological sex within the cohort, the potential cohort enrichment of higher SES participants, and the inability to correct for smoking-related effects. Additionally, in analyzing grey matter volume of fronto-limbic brain regions as outcomes of interest, we have omitted surface area- or cortical thickness-specific effects. The current study also falls short in establishing whether the mediation by peripheral 5mC modules is causal in nature. Longitudinal data could provide more precise insight into whether such relationships exist. Future studies on this topic should capture longitudinal data from a diverse, increased sample size and could investigate genetic factors or tissues of etiological interest.

## Conclusions

The current study showed that, in support of prior literature, exposure to poor childhood FEH is associated with low fronto-limbic BRV as measured in young adulthood. Newly reported here is the finding that saliva-derived 5mC modules mediate the FEH and BRV relationship and are enriched for immune system, CNS-related, and metabolic functions; with additional validation in independent cohorts, these 5mC modules could potentially be used as peripheral biomarkers of poor FEH exposure during childhood. Overall, the findings of the current study support the neuroimmune network hypothesis[6], extend the body of work highlighting neurodevelopmental variability associated with childhood ASE exposure, and inform a potential molecular mechanism of biologic embedding. Future research on these peripheral signatures could validate their use as proxies/biomarkers of perturbed underlying neurobiology in response to poor FEH exposure and could inform further investigation into primarily effected tissue such as endocrine, immune, and CNS cell types.

## Materials and methods

### Participants

The current study draws on data from 98 university-age students (19.8±1.2 years old; 69% women; 49% white) who successfully completed the Duke Neurogenetics Study (DNS). The DNS aims to assess the associations among a wide range of behavioral, neural, and genetic variables in a large sample of young adults, with one of the core goals being to establish a link between these various phenotypes and psychopathology. This study was approved by the Duke University Medical Center Institutional Review Board, and all experiments were performed in accordance to its guidelines. Prior to the study, all participants provided informed consent. To be eligible for DNS, all participants were free of: 1) medical diagnoses of cancer, stroke, head injury with loss of consciousness, untreated migraine headaches, diabetes requiring insulin treatment, chronic kidney, or liver disease; 2) use of psychotropic, glucocorticoid, or hypolipidemic medication; and 3) conditions affecting cerebral blood flow and metabolism (e.g., hypertension)[75].

### Family emotional health (FEH)

Participants were asked to complete the Family History Questionnaire (FHQ), which produced the current study’s measure of FEH. The FHQ is composed *fully* of questions from previously validated inventories [76–82]; fifty-five out of 70 questions were included from the Family History Screen (FHS)[76, 77]. The FHQ and FHS both capture family-wide psychiatric illness, but the FHQ is more encompassing of other ASEs, including cognitive decline of family members[78], externalizing behaviors[79], exposure to smoking[80], and drug/alcohol abuse treatment[81, 82]. The summed responses from 70 “yes/no” questions based on the aforementioned topics from the FHQ represent the current study’s measure of FEH (Supplementary Table 1). Each “no” response corresponded to an additional score of one, with lower values representing poor FEH.

### Cumulative perceived impact of past year stressful life events (past year SLEs)

Participants were administered an inventory measuring the cumulative perceived impact of SLEs from the past year (“past year SLEs”). Prior research reported associations between stress exposure and significant variability in fronto-limbic brain region volumes (BRVs)[17–19]. Therefore, throughout the current study, we controlled for the effect of past years SLEs using a summation of 45 negatively valenced items[83, 84] from the Life Events Scale for Students[85].

### Neuroimaging

ASEs and exposures similar to poor FEH are known to impact fronto-limbic pathways in the human CNS[17–23]. In addition, a meta-analysis has shown that both the hippocampus and amygdala have hemisphere-specific volume differences in healthy adults[86], and ASEs are known to have hemisphere-specific effects on fronto-limbic brain regions[23, 87]. Therefore, hemisphere-specific amygdala, hippocampus dorso-lateral prefrontal cortex (dlPFC), and medial PFC (mPFC) volume measures were estimated. Volume measurements of dlPFC and mPFC were chosen as outcome variables from the frontal cortex due to the opposing nature of their afferent and efferent projections to hippocampus and amygdala, and their functional relationships with each region[88]. Participants were scanned on one of two identical research-dedicated GE MR750 3T scanners at the Duke-UNC Brain Imaging and Analysis Center, and measures were collected, pre-processed, and finalized in accordance with previously published methods[89]. Briefly, anatomical images for each subject were skull-stripped, intensity-normalized, and mapped to a study-specific average template. Region-specific border definitions were made using the Desikan-Killiany-Tourville scheme[90].

### Molecular

Saliva was collected from participants using the Oragene-DNA OG-500 kit (Oragene; Ottawa, Canada). DNA was extracted and cleaned using the DNA Genotek prepIT PT-L2P kit (DNA Genotek Inc; Ottawa, Canada) using manufacturer recommended methods. Purity of extracted DNA samples was assessed by absorbance using Nanodrop 1000 spectrophotometer (Thermo Fisher Scientific Inc; Waltham, Massachusetts). The quantity of double-stranded DNA was assessed using Quant-iT PicoGreen dsDNA kits with manufacturer recommended protocols (Invitrogen; Carlsbad, California). A total of 500 ng of genomic DNA was bisulfite-converted (BSC) using manufacturer-recommended EZ DNAm kits (Zymo Research; Irvine, California). After conversion, BSC DNA was applied to the Infinium MethylationEPIC BeadChip (Illumina; San Diego, California) (850k) to measure 5mC at ~850k loci.

### 5mC pre-processing

Beta-values measured from the 850k platform were background corrected in GenomeStudio, quality controlled, and filtered according to previously published methods[91]. All quality control and pre-processing was performed in R, version 3.6.1[92]. These steps removed low quality and potentially cross-hybridizing probes, quantile-normalized probe beta-values, and removed technical and batch effects[93–96]. 5mC beta-values were variance stabilized and logit-transformed into M-values[97]. X- and Y-chromosome-mapped probes were removed, along with *rs*-mapped probes. The remaining ~739k probes were then subset to include only those with observed nominally significant Pearson correlation (p<0.05) between saliva and brain tissue from the ImageCpG data repository[98]. This was done to focus the analysis on loci with greater prospect for proxy or surrogate status with etiologically relevant CNS tissue. Afterwards, 62,422 probes remained.

### Cellular heterogeneity

Cell heterogeneity was estimated using a reference-free deconvolution method[99, 100]. Briefly, the top 15k most variable CpG sites were selected from the pre-processed/quality controlled 850k data and used to estimate the number of cell types and generate a matrix containing the proportions. Based on these methods, the number of cell types was set at five. Estimated proportions were used as covariates in relevant analyses to account for cellular heterogeneity.

### Genomic ancestry

To avoid potential inaccuracies and confounding effects of self-reported race/ethnicity, genetic ancestry was modeled using multi-dimensional scaling (MDS) measures extracted from participant genomic data using PLINK[101]. Using previously collected GWAS data from the DNS, the first four MDS genetic ancestry measures were calculated and used as covariates across pertinent models based on visual inspection of scree plots. This methodology is in line with previous publications[75].

### Probe clustering

In order to remove non-desired effects, we fit linear models with age, validated biological sex, cellular heterogeneity, and genomic ancestry as predictors of probe-wise 5mC M-value. For each probe, residual values (“residualized M-values”) were extracted for clustering. Taking the 62,422 residualized M-values, the “WGCNA” R package was used to build a co-methylation network[102]. First, scale-free topology model fit was analyzed. As recommended, a soft-threshold value of four was chosen based on the lowest power at which adjusted R^2^>0.90. Adjacency and dissimilarity matrices were generated, and unsupervised hierarchical clustering was used to generate a clustered, residual M-value network. Setting a minimum cluster size of 10 generated 194 modules, identified by a unique color, for which module eigengenes (MEs) were calculated.

### Statistical analyses

In order to understand the relationships between variables, we computed Pearson correlations and mapped their correlation coefficients. Based on these correlations, we conducted a set of analyses, as shown in Figure 1. In Arm A analyses, FEH was used as a predictor of hemisphere-specific BRVs, while including age, biological sex, four genomic ancestry MDS measures, past year SLEs, and TBV as covariates. In Arm B analyses, FEH was used as a predictor of ME values, while including past year SLEs as a covariate. Age, sex, and genomic ancestry effects were accounted for previously by using residualized M-values as input for clustering. In Arm C analyses, ME values were used as individual predictors of BRV, while including the same covariates as in Arm A. Throughout the current research, past year SLEs were included as a covariate because our FEH measure only captures SLEs from childhood, and recent stress exposure is associated with variability in our outcome variables[17, 18, 23, 103]. TBV was included as a covariate but, where pertinent, non-TBV controlled model results are also reported. Within each phase of the analyses, non-standardized continuous measures were used resulting in nonstandardized effect estimates. In addition, nominal p-values were corrected for multiple hypothesis testing by controlling the false discovery rate (FDR=0.10) using the Benjamini Hochberg (BH) procedure[104]. Briefly, for each nominal p-value, a BH critical value was calculated where nominal p-value’s assigned rank over the number of tests was multiplied by the accepted FDR. Nominal p-values less than this threshold were deemed BH-significant. Due to the exploratory nature of the current work, both nominal and BH-significant terms were considered for interpretation.

### Mediation analyses

To investigate whether the effect of poor FEH on hemisphere-specific BRV is *statistically* mediated via peripheral 5mC signatures, MEs were tested for mediating status between FEH and hemisphere-specific BRVs using the “mediation” package in R[105] (Figure 1). Importantly, only hemisphere-specific BRVs associated with FEH (Figure 1, Arm A) were considered. Similarly, MEs tested for mediation included *only those* associated with both FEH (Figure 1, Arm B) and hemisphere-specific BRV (Figure 1, Arm C). Mediation model inputs were assembled per recommended “mediation” package protocol. Therefore, Arm A (plus ME as a covariate) and Arm B models were used as inputs. For each ME, indirect effects (IDE), direct effects (DE), and total effects (TE) were calculated as a result of 10,000 non-parametric bootstrap simulations. Consistent with published methods[23], we considered an ME a full mediator if the DE=0 while the IDE and TE≠ 0, or a partial mediator if the DE, IDE, and TE≠ 0. Individual probes from full mediator modules were assessed for mediation status as well.

### Gene set enrichment

To assess the underlying methylomic network enrichment of the ~62,000 brain-saliva correlated probes, individual residualized probe M-values were used as predictors of FEH in Bayesian regression models. Age, sex, genomic ancestry measures, cell heterogeneity measures, and past year SLEs were included as covariates. From this analysis, BH-significant probe p-values were extracted and used as input to gene set enrichment analyses (GSEA) in the “methylGSA” package[106]. GO sets composed of 50 to 1,000 genes were allowed, which eliminated high-level GO-terms such as “biological process” and facilitated testing of 3,186 GO sets. To produce a condensed summary of non-redundant GO-terms, the web-based tool “Revigo” was used [107].

## Declarations

### Ethics approval and consent to participate

This study was approved by the Duke University Medical Center Institutional Review Board, and all experiments were performed in accordance to its guidelines.

### Consent for publication

Not applicable.

### Availability of data and materials

The ImageCpG dataset supporting the conclusions of this article is available at Gene Expression Omnibus (GEO) Accession GSE111165; http://han-lab.org/methylation/default/imageCpG#. The DNS 850K datasets used and/or analyzed during the current study are available from the corresponding author on reasonable request.

### Authors’ contributions

JRP was a major contributor in study design, methodology, statistical analyses, and in writing the manuscript. ACB was a major contributor in methodology and writing the manuscript. GSK was a major contributor in methodology and writing the manuscript. DA was a major contributor in methodology and writing the manuscript. ARK was a major contributor in writing the manuscript. KCK was a major contributor in writing the manuscript. ARH was a major contributor in designing methodology and writing the manuscript. MU was a major contributor in study design, methodology, and in writing the manuscript. All authors read and approved the final manuscript.

## Acknowledgements

We thank the Duke Neurogenetics Study participants and the staff of the Laboratory of NeuroGenetics.

## References

1. Stambaugh LF, Forman-Hoffman V, Williams J, Pemberton MR, Ringeisen H, Hedden SL, et al. Prevalence of serious mental illness among parents in the United States: results from the National Survey of Drug Use and Health, 2008-2014. Ann Epidemiol. 2017;27:222–4.

2. National Research Council (US) and Institute of Medicine (US) Committee on Depression PP, England MJ, Sim LJ. Associations Between Depression in Parents and Parenting, Child Health, and Child Psychological Functioning. National Academies Press (US); 2009. https://www.ncbi.nlm.nih.gov/books/NBK215128/. Accessed 13 May 2020.

3. Dean K, Stevens H, Mortensen PB, Murray RM, Walsh E, Pedersen CB. Full Spectrum of Psychiatric Outcomes Among Offspring With Parental History of Mental Disorder. Arch Gen Psychiatry. 2010;67:822–9.

4. Connell AM, Goodman SH. The association between psychopathology in fathers versus mothers and children’s internalizing and externalizing behavior problems: a meta-analysis. Psychol Bull. 2002;128:746–73.

5. Suveg C, Shaffer A, Morelen D, Thomassin K. Links Between Maternal and Child Psychopathology Symptoms: Mediation Through Child Emotion Regulation and Moderation Through Maternal Behavior. Child Psychiatry Hum Dev. 2011;42:507.

6. Nusslock R, Miller GE. Early-Life Adversity and Physical and Emotional Health Across the Lifespan: A Neuroimmune Network Hypothesis. Biol Psychiatry. 2016;80:23–32.

7. Dunn EC, Soare TW, Zhu Y, Simpkin AJ, Suderman MJ, Klengel T, et al. Sensitive Periods for the Effect of Childhood Adversity on DNA Methylation: Results From a Prospective, Longitudinal Study. Biol Psychiatry. 2019;85:838–49.

8. Walter KH, Palmieri PA, Gunstad J. More than symptom reduction: Changes in executive function over the course of PTSD treatment. J Trauma Stress. 2010;23:292–5.

9. Misiak B, Beszłej JA, Kotowicz K, Szewczuk-Bogusławska M, Samochowiec J, Kucharska-Mazur J, et al. Cytokine alterations and cognitive impairment in major depressive disorder: From putative mechanisms to novel treatment targets. Prog Neuropsychopharmacol Biol Psychiatry. 2018;80:177–88.

10. Marganska A, Gallagher M, Miranda R. Adult attachment, emotion dysregulation, and symptoms of depression and generalized anxiety disorder. Am J Orthopsychiatry. 2013;83:131–41.

11. Green MF. Cognitive impairment and functional outcome in schizophrenia and bipolar disorder. J Clin Psychiatry. 2006;67:e12.

12. Anisman H. Cascading effects of stressors and inflammatory immune system activation: implications for major depressive disorder. J Psychiatry Neurosci JPN. 2009;34:4–20.

13. Cohen S, Tyrrell DA, Smith AP. Negative life events, perceived stress, negative affect, and susceptibility to the common cold. J Pers Soc Psychol. 1993;64:131–40.

14. Segerstrom SC, Miller GE. Psychological stress and the human immune system: a meta-analytic study of 30 years of inquiry. Psychol Bull. 2004;130:601–30.

15. Salleh MohdR. Life Event, Stress and Illness. Malays J Med Sci MJMS. 2008;15:9–18.

16. Lupien SJ, King S, Meaney MJ, McEwen BS. Child’s stress hormone levels correlate with mother’s socioeconomic status and depressive state. Biol Psychiatry. 2000;48:976–80.

17. Pagliaccio D, Luby JL, Bogdan R, Agrawal A, Gaffrey MS, Belden AC, et al. Stress-System Genes and Life Stress Predict Cortisol Levels and Amygdala and Hippocampal Volumes in Children. Neuropsychopharmacology. 2014;39:1245–53.

18. Hanson JL, Nacewicz BM, Sutterer MJ, Cayo AA, Schaefer SM, Rudolph KD, et al. Behavior Problems After Early Life Stress: Contributions of the Hippocampus and Amygdala. Biol Psychiatry. 2015;77:314–23.

19. Kim P, Evans GW, Angstadt M, Ho SS, Sripada CS, Swain JE, et al. Effects of childhood poverty and chronic stress on emotion regulatory brain function in adulthood. Proc Natl Acad Sci U S A. 2013;110:18442–7.

20. Swartz JR, Knodt AR, Radtke SR, Hariri AR. A Neural Biomarker of Psychological Vulnerability to Future Life Stress. Neuron. 2015;85:505–11.

21. Swartz JR, Hariri AR, Williamson DE. An epigenetic mechanism links socioeconomic status to changes in depression-related brain function in high-risk adolescents. Mol Psychiatry. 2017;22:209–14.

22. Luby JL, Barch DM, Belden A, Gaffrey MS, Tillman R, Babb C, et al. Maternal support in early childhood predicts larger hippocampal volumes at school age. Proc Natl Acad Sci. 2012;109:2854–9.

23. Luby J, Belden A, Botteron K, Marrus N, Harms MP, Babb C, et al. The effects of poverty on childhood brain development: the mediating effect of caregiving and stressful life events. JAMA Pediatr. 2013;167:1135–42.

24. Hasler G, Northoff G. Discovering imaging endophenotypes for major depression. Mol Psychiatry. 2011;16:604–19.

25. Manoach DS, Agam Y. Neural markers of errors as endophenotypes in neuropsychiatric disorders. Front Hum Neurosci. 2013;7. doi:10.3389/fnhum.2013.00350.

26. Matsubara T, Matsuo K, Harada K, Nakano M, Nakashima M, Watanuki T, et al. Distinct and Shared Endophenotypes of Neural Substrates in Bipolar and Major Depressive Disorders. PLOS ONE. 2016;11:e0168493.

27. Toyokawa S, Uddin M, Koenen KC, Galea S. How does the social environment ‘get into the mind’? Epigenetics at the intersection of social and psychiatric epidemiology. Soc Sci Med 1982. 2012;74:67–74.

28. Klengel T, Mehta D, Anacker C, Rex-Haffner M, Pruessner JC, Pariante CM, et al. Allele-specific FKBP5 DNA demethylation mediates gene–childhood trauma interactions. Nat Neurosci. 2013;16:33–41.

29. Weaver ICG. Integrating Early Life Experience, Gene Expression, Brain Development, and Emergent Phenotypes. In: Advances in Genetics. Elsevier; 2014. p. 277–307. doi:10.1016/B978-0-12-800222-3.00011-5.

30. Jaenisch R, Bird A. Epigenetic regulation of gene expression: how the genome integrates intrinsic and environmental signals. Nat Genet. 2003;33 Suppl:245–54.

31. Zilberman D, Gehring M, Tran RK, Ballinger T, Henikoff S. Genome-wide analysis of Arabidopsis thaliana DNA methylation uncovers an interdependence between methylation and transcription. Nat Genet. 2007;39:61–9.

32. Talens RP, Boomsma DI, Tobi EW, Kremer D, Jukema JW, Willemsen G, et al. Variation, patterns, and temporal stability of DNA methylation: considerations for epigenetic epidemiology. FASEB J Off Publ Fed Am Soc Exp Biol. 2010;24:3135–44.

33. Michaud EJ, van Vugt MJ, Bultman SJ, Sweet HO, Davisson MT, Woychik RP. Differential expression of a new dominant agouti allele (Aiapy) is correlated with methylation state and is influenced by parental lineage. Genes Dev. 1994;8:1463–72.

34. Smith AK, Kilaru V, Klengel T, Mercer KB, Bradley B, Conneely KN, et al. DNA extracted from saliva for methylation studies of psychiatric traits: evidence tissue specificity and relatedness to brain. Am J Med Genet Part B Neuropsychiatr Genet Off Publ Int Soc Psychiatr Genet. 2015;168B:36–44.

35. Perroud N, Paoloni-Giacobino A, Prada P, Olié E, Salzmann A, Nicastro R, et al. Increased methylation of glucocorticoid receptor gene (NR3C1) in adults with a history of childhood maltreatment: a link with the severity and type of trauma. Transl Psychiatry. 2011;1:e59.

36. Tyrka AR, Price LH, Marsit C, Walters OC, Carpenter LL. Childhood adversity and epigenetic modulation of the leukocyte glucocorticoid receptor: preliminary findings in healthy adults. PloS One. 2012;7:e30148.

37. Uddin M, Aiello AE, Wildman DE, Koenen KC, Pawelec G, de los Santos R, et al. Epigenetic and immune function profiles associated with posttraumatic stress disorder. Proc Natl Acad Sci U S A. 2010;107:9470–5.

38. Smith AK, Conneely KN, Kilaru V, Mercer KB, Weiss TE, Bradley B, et al. Differential immune system DNA methylation and cytokine regulation in post-traumatic stress disorder. Am J Med Genet Part B Neuropsychiatr Genet Off Publ Int Soc Psychiatr Genet. 2011;156B:700–8.

39. Davies MN, Volta M, Pidsley R, Lunnon K, Dixit A, Lovestone S, et al. Functional annotation of the human brain methylome identifies tissue-specific epigenetic variation across brain and blood. Genome Biol. 2012;13:R43.

40. Hannon E, Lunnon K, Schalkwyk L, Mill J. Interindividual methylomic variation across blood, cortex, and cerebellum: implications for epigenetic studies of neurological and neuropsychiatric phenotypes. Epigenetics. 2015;10:1024–32.

41. Yamamoto T, Toki S, Siegle GJ, Takamura M, Takaishi Y, Yoshimura S, et al. Increased amygdala reactivity following early life stress: a potential resilience enhancer role. BMC Psychiatry. 2017;17. doi:10.1186/s12888-017-1201-x.

42. Booij L, Szyf M, Carballedo A, Frey E-M, Morris D, Dymov S, et al. DNA methylation of the serotonin transporter gene in peripheral cells and stress-related changes in hippocampal volume: a study in depressed patients and healthy controls. PloS One. 2015;10:e0119061.

43. Frodl T, Szyf M, Carballedo A, Ly V, Dymov S, Vaisheva F, et al. DNA methylation of the serotonin transporter gene (SLC6A4) is associated with brain function involved in processing emotional stimuli. J Psychiatry Neurosci JPN. 2015;40:296–305.

44. Ismaylova E, Lévesque ML, Pomares FB, Szyf M, Nemoda Z, Fahim C, et al. Serotonin transporter promoter methylation in peripheral cells and neural responses to negative stimuli: A study of adolescent monozygotic twins. Transl Psychiatry. 2018;8:1–9.

45. Vukojevic V, Kolassa I-T, Fastenrath M, Gschwind L, Spalek K, Milnik A, et al. Epigenetic modification of the glucocorticoid receptor gene is linked to traumatic memory and post-traumatic stress disorder risk in genocide survivors. J Neurosci Off J Soc Neurosci. 2014;34:10274–84.

46. Schechter DS, Moser DA, Paoloni-Giacobino A, Stenz L, Gex-Fabry M, Aue T, et al. Methylation of NR3C1 is related to maternal PTSD, parenting stress and maternal medial prefrontal cortical activity in response to child separation among mothers with histories of violence exposure. Front Psychol. 2015;6:690.

47. Tozzi L, Farrell C, Booij L, Doolin K, Nemoda Z, Szyf M, et al. Epigenetic Changes of FKBP5 as a Link Connecting Genetic and Environmental Risk Factors with Structural and Functional Brain Changes in Major Depression. Neuropsychopharmacol Off Publ Am Coll Neuropsychopharmacol. 2018;43:1138–45.

48. Sadeh N, Wolf EJ, Logue MW, Hayes JP, Stone A, Griffin LM, et al. EPIGENETIC VARIATION AT SKA2 PREDICTS SUICIDE PHENOTYPES AND INTERNALIZING PSYCHOPATHOLOGY. Depress Anxiety. 2016;33:308–15.

49. Sadeh N, Spielberg JM, Logue MW, Wolf EJ, Smith AK, Lusk J, et al. SKA2 methylation is associated with decreased prefrontal cortical thickness and greater PTSD severity among trauma-exposed veterans. Mol Psychiatry. 2016;21:357–63.

50. Schaafsma S m, Riedstra B j, Pfannkuche K a, Bouma A, Groothuis T g. g. Epigenesis of behavioural lateralization in humans and other animals. Philos Trans R Soc B Biol Sci. 2009;364:915–27.

51. Gotts SJ, Jo HJ, Wallace GL, Saad ZS, Cox RW, Martin A. Two distinct forms of functional lateralization in the human brain. Proc Natl Acad Sci. 2013;110:E3435–44.

52. Francks C. Exploring human brain lateralization with molecular genetics and genomics. Ann N Y Acad Sci. 2015;1359:1–13.

53. Kim E, Garrett A, Boucher S, Park M-H, Howe M, Sanders E, et al. Inhibited Temperament and Hippocampal Volume in Offspring of Parents with Bipolar Disorder. J Child Adolesc Psychopharmacol. 2016;27:258–65.

54. Haren NEM van, Setiaman N, Koevoets MGJC, Baalbergen H, Kahn RS, Hillegers MHJ. Brain structure, IQ, and psychopathology in young offspring of patients with schizophrenia or bipolar disorder. Eur Psychiatry. 2020;63. doi:10.1192/j.eurpsy.2019.19.

55. Klengel T, Pape J, Binder EB, Mehta D. The role of DNA methylation in stress-related psychiatric disorders. Neuropharmacology. 2014;80:115–32.

56. Tomoda A, Suzuki H, Rabi K, Sheu Y-S, Polcari A, Teicher MH. Reduced Prefrontal Cortical Gray Matter Volume in Young Adults Exposed to Harsh Corporal Punishment. NeuroImage. 2009;47 Suppl 2:T66–71.

57. Carrion VG, Weems CF, Richert K, Hoffman BC, Reiss AL. Decreased Prefrontal Cortical Volume Associated With Increased Bedtime Cortisol in Traumatized Youth. Biol Psychiatry. 2010;68:491–3.

58. Corbo V, Salat DH, Amick MM, Leritz EC, Milberg WP, McGlinchey RE. Reduced cortical thickness in veterans exposed to early life trauma. Psychiatry Res. 2014;223:53–60.

59. Kent WJ, Sugnet CW, Furey TS, Roskin KM, Pringle TH, Zahler AM, et al. The human genome browser at UCSC. Genome Res. 2002;12:996–1006.

60. Rosenbloom KR, Sloan CA, Malladi VS, Dreszer TR, Learned K, Kirkup VM, et al. ENCODE Data in the UCSC Genome Browser: year 5 update. Nucleic Acids Res. 2013;41:D56–63.

61. Tian G, Thomas S, Cowan NJ. Effect of TBCD and its regulatory interactor Arl2 on tubulin and microtubule integrity. Cytoskelet Hoboken NJ. 2010;67:706–14.

62. Flex E, Niceta M, Cecchetti S, Thiffault I, Au MG, Capuano A, et al. Biallelic Mutations in TBCD, Encoding the Tubulin Folding Cofactor D, Perturb Microtubule Dynamics and Cause Early-Onset Encephalopathy. Am J Hum Genet. 2016;99:962–73.

63. Miyake N, Fukai R, Ohba C, Chihara T, Miura M, Shimizu H, et al. Biallelic TBCD Mutations Cause Early-Onset Neurodegenerative Encephalopathy. Am J Hum Genet. 2016;99:950–61.

64. Guo S, Zhu Q, Jiang T, Wang R, Shen Y, Zhu X, et al. Genome-wide DNA methylation patterns in CD4+ T cells from Chinese Han patients with rheumatoid arthritis. Mod Rheumatol. 2017;27:441–7.

65. Song H, Fang F, Tomasson G, Arnberg FK, Mataix-Cols D, Cruz LF de la, et al. Association of Stress-Related Disorders With Subsequent Autoimmune Disease. JAMA. 2018;319:2388–400.

66. Duclot F, Kabbaj M. The Role of Early Growth Response 1 (EGR1) in Brain Plasticity and Neuropsychiatric Disorders. Front Behav Neurosci. 2017;11. doi:10.3389/fnbeh.2017.00035.

67. Zhang W, Liu HT. MAPK signal pathways in the regulation of cell proliferation in mammalian cells. Cell Res. 2002;12:9–18.

68. Wefers B, Hitz C, Hölter SM, Trümbach D, Hansen J, Weber P, et al. MAPK Signaling Determines Anxiety in the Juvenile Mouse Brain but Depression-Like Behavior in Adults. PLOS ONE. 2012;7:e35035.

69. Bruchas MR, Schindler AG, Shankar H, Messinger DI, Miyatake M, Land BB, et al. Selective p38α MAPK Deletion in Serotonergic Neurons Produces Stress Resilience in Models of Depression and Addiction. Neuron. 2011;71:498–511.

70. Zhu C-B, Lindler KM, Owens AW, Daws LC, Blakely RD, Hewlett WA. Interleukin-1 Receptor Activation by Systemic Lipopolysaccharide Induces Behavioral Despair Linked to MAPK Regulation of CNS Serotonin Transporters. Neuropsychopharmacology. 2010;35:2510–20.

71. Firestein GS. Immunologic mechanisms in the pathogenesis of rheumatoid arthritis. J Clin Rheumatol Pract Rep Rheum Musculoskelet Dis. 2005;11 3 Suppl:S39–44.

72. Fifield J, Tennen H, Reisine S, McQuillan J. Depression and the long-term risk of pain, fatigue, and disability in patients with rheumatoid arthritis. Arthritis Rheum. 1998;41:1851–7.

73. Malemud CJ, Miller AH. Pro-inflammatory cytokine-induced SAPK/MAPK and JAK/STAT in rheumatoid arthritis and the new anti-depression drugs. Expert Opin Ther Targets. 2008;12:171–83.

74. Chan KL, Cathomas F, Russo SJ. Central and Peripheral Inflammation Link Metabolic Syndrome and Major Depressive Disorder. Physiology. 2019;34:123–33.

75. Kim MJ, Avinun R, Knodt AR, Radtke SR, Hariri AR. Neurogenetic plasticity and sex influence the link between corticolimbic structural connectivity and trait anxiety. Sci Rep. 2017;7:10959.

76. Weissman MM, Wickramaratne P, Adams P, Wolk S, Verdeli H, Olfson M. Brief screening for family psychiatric history: the family history screen. Arch Gen Psychiatry. 2000;57:675–82.

77. Milne BJ, Caspi A, Crump R, Poulton R, Rutter M, Sears MR, et al. The Validity of the Family History Screen for Assessing Family History of Mental Disorders. Am J Med Genet Part B Neuropsychiatr Genet Off Publ Int Soc Psychiatr Genet. 2009;0:41–9.

78. Devi G, Marder K, Schofield PW, Tang MX, Stern Y, Mayeux R. Validity of family history for the diagnosis of dementia among siblings of patients with late-onset Alzheimer’s disease. Genet Epidemiol. 1998;15:215–23.

79. Bidaut-Russell M, Reich W, Cottler LB, Robins LN, Compton WM, Mattison RE. The Diagnostic Interview Schedule for Children (PC-DISC v.3.0): parents and adolescents suggest reasons for expecting discrepant answers. J Abnorm Child Psychol. 1995;23:641–59.

80. Williams RR, Hunt SC, Barlow GK, Chamberlain RM, Weinberg AD, Cooper HP, et al. Health family trees: a tool for finding and helping young family members of coronary and cancer prone pedigrees in Texas and Utah. Am J Public Health. 1988;78:1283–6.

81. Skinner HA. The drug abuse screening test. Addict Behav. 1982;7:363–71.

82. Selzer ML, Vinokur A, van Rooijen L. A self-administered Short Michigan Alcoholism Screening Test (SMAST). J Stud Alcohol. 1975;36:117–26.

83. Avinun R, Nevo A, Knodt AR, Elliott ML, Radtke SR, Brigidi BD, et al. Reward-Related Ventral Striatum Activity Buffers against the Experience of Depressive Symptoms Associated with Sleep Disturbances. J Neurosci Off J Soc Neurosci. 2017;37:9724–9.

84. Nikolova YS, Bogdan R, Brigidi BD, Hariri AR. Ventral Striatum Reactivity to Reward and Recent Life Stress Interact to Predict Positive Affect. Biol Psychiatry. 2012;72:157–63.

85. Clements K, Turpin G. The Life Events Scale for Students: Validation for use with British samples. Personal Individ Differ. 1996;20:747–51.

86. Pedraza O, Bowers D, Gilmore R. Asymmetry of the hippocampus and amygdala in MRI volumetric measurements of normal adults. J Int Neuropsychol Soc. 2004;10:664–78.

87. Lindauer RJL, Vlieger E-J, Jalink M, Olff M, Carlier IVE, Majoie CBLM, et al. Smaller hippocampal volume in Dutch police officers with posttraumatic stress disorder. Biol Psychiatry. 2004;56:356–63.

88. Kovner R, Oler JA, Kalin NH. Cortico-Limbic Interactions Mediate Adaptive and Maladaptive Responses Relevant to Psychopathology. Am J Psychiatry. 2019;176:987–99.

89. Elsayed NM, Kim MJ, Fields KM, Olvera RL, Hariri AR, Williamson DE. Trajectories of Alcohol Initiation and Use During Adolescence: The Role of Stress and Amygdala Reactivity. J Am Acad Child Adolesc Psychiatry. 2018;57:550–60.

90. Klein A, Tourville J. 101 Labeled Brain Images and a Consistent Human Cortical Labeling Protocol. Front Neurosci. 2012;6. doi:10.3389/fnins.2012.00171.

91. Ratanatharathorn A, Boks MP, Maihofer AX, Aiello AE, Amstadter AB, Ashley-Koch AE, et al. Epigenome-wide association of PTSD from heterogeneous cohorts with a common multi-site analysis pipeline. Am J Med Genet Part B Neuropsychiatr Genet Off Publ Int Soc Psychiatr Genet. 2017;174:619–30.

92. R Core Team (2019). R: A language and environment for statistical computing. R Foundation for Statistical Computing, Vienna, Austria. URL https://www.R-project.org/.

93. Barfield RT, Kilaru V, Smith AK, Conneely KN. CpGassoc: an R function for analysis of DNA methylation microarray data. Bioinforma Oxf Engl. 2012;28:1280–1.

94. McCartney DL, Walker RM, Morris SW, McIntosh AM, Porteous DJ, Evans KL. Identification of polymorphic and off-target probe binding sites on the Illumina Infinium MethylationEPIC BeadChip. Genomics Data. 2016;9:22–4.

95. Teschendorff AE, Marabita F, Lechner M, Bartlett T, Tegner J, Gomez-Cabrero D, et al. A beta-mixture quantile normalization method for correcting probe design bias in Illumina Infinium 450 k DNA methylation data. Bioinforma Oxf Engl. 2013;29:189–96.

96. Pidsley R, Y Wong CC, Volta M, Lunnon K, Mill J, Schalkwyk LC. A data-driven approach to preprocessing Illumina 450K methylation array data. BMC Genomics. 2013;14:293.

97. Du P, Zhang X, Huang C-C, Jafari N, Kibbe WA, Hou L, et al. Comparison of Beta-value and M-value methods for quantifying methylation levels by microarray analysis. BMC Bioinformatics. 2010;11:587.

98. Braun PR, Han S, Hing B, Nagahama Y, Gaul LN, Heinzman JT, et al. Genome-wide DNA methylation comparison between live human brain and peripheral tissues within individuals. Transl Psychiatry. 2019;9:1–10.

99. Houseman EA, Molitor J, Marsit CJ. Reference-free cell mixture adjustments in analysis of DNA methylation data. Bioinforma Oxf Engl. 2014;30:1431–9.

100. Houseman EA, Kile ML, Christiani DC, Ince TA, Kelsey KT, Marsit CJ. Reference-free deconvolution of DNA methylation data and mediation by cell composition effects. BMC Bioinformatics. 2016;17:259.

101. Purcell S, Neale B, Todd-Brown K, Thomas L, Ferreira MAR, Bender D, et al. PLINK: a tool set for wholegenome association and population-based linkage analyses. Am J Hum Genet. 2007;81:559–75.

102. Langfelder P, Horvath S. WGCNA: an R package for weighted correlation network analysis. BMC Bioinformatics. 2008;9:559.

103. Unternaehrer E, Luers P, Mill J, Dempster E, Meyer AH, Staehli S, et al. Dynamic changes in DNA methylation of stress-associated genes (OXTR, BDNF) after acute psychosocial stress. Transl Psychiatry. 2012;2:e150.

104. Benjamini Y, Hochberg Y. Controlling the False Discovery Rate: A Practical and Powerful Approach to Multiple Testing. J R Stat Soc Ser B Methodol. 1995;57:289–300.

105. Tingley D, Yamamoto T, Hirose K, Keele L, Imai K. mediation: R Package for Causal Mediation Analysis. J Stat Softw. 2014;59. doi:10.18637/jss.v059.i05.

106. Ren X, Kuan PF. methylGSA: a Bioconductor package and Shiny app for DNA methylation data length bias adjustment in gene set testing. Bioinforma Oxf Engl. 2019;35:1958–9.

107. Supek F, Bošnjak M, Škunca N, Šmuc T. REVIGO summarizes and visualizes long lists of gene ontology terms. PloS One. 2011;6:e21800.

